# Immune-cell depleted diffuse large B-cell lymphomas have reduced expression of MHC class I

**DOI:** 10.64898/2026.09.01.748487

**Authors:** Kanutte Huse, Yngvild N. Blaker, Jillian F. Wise, Marie Hairing Enemark, Maare Arffman, Ankush Sharma, Leo Meriranta, Sigve Nakken, Kathrine Isaksen, Ivana Spasevska, Vera Hilden, Chloé B. Steen, David Warren, Bent Honoré, Klaus Beiske, Erlend B. Smeland, Qiang Pan-Hammarström, Maja Ludvigsen, Sirpa Leppä, Harald Holte, June H. Myklebust

## Abstract

Immunotherapy has transformed treatment for many cancers. In the aggressive and genetically heterogeneous diffuse large B-cell lymphoma (DLBCL), CD19 CAR T-cell therapy is highly effective, whereas immune checkpoint blockade has shown limited benefit. Loss of MHC expression is a common mechanism to escape T-cell cytotoxicity, and loss of MHC class I (MHC-I) and II are frequent in DLBCL. We applied imaging mass cytometry to diagnostic biopsies from younger, high-risk DLBCL patients to map the tumor microenvironment (TME) spatial architecture in relation to tumor cell MHC expression, mutational status, transcriptomic and proteomic profiles. Neighborhood analyses identified four TME subtypes: immune-cell depleted and three immune-infiltrated types (mixed, CD4 T cell-rich, CD8 T-cell/macrophage-rich). Depleted cases had shorter overall survival (*p* = 0.033) and increased expression of proteins involved in DNA replication and proliferation markers compared to infiltrated cases. Tumor cell MHC-I expression was heterogeneous. Cases with low frequency of MHC-I^pos^ tumor cells were enriched for the depleted TME type. MHC-I^pos^ tumor cells were surrounded by CD4 and CD8 T cells and M1 macrophages, whereas MHC-I^neg^ tumor cells were closer to other MHC-I^neg^ tumor cells. These findings suggest that TME-based classification incorporating tumor cell MHC-I status may improve individualized immunotherapy selection.

## Introduction

Immunotherapy has changed the therapeutic landscape for different types of cancer. For diffuse large B-cell lymphoma (DLBCL), the most common form of aggressive B-cell non-Hodgkin lymphoma, over 60% of patients are cured by standard chemo-immunotherapy^1^. Previously, relapsed or refractory DLBCL had very poor outcomes, but immunotherapy with CD19 CAR T cells has now induced long-term disease control in about 40% of the patients^2,3^. Bispecific T-cell engagers represent a promising off-the shelf alternative^4,5^. In contrast, treatment with immune checkpoint blockade monotherapy had limited efficacy in DLBCL^6^.

DLBCL tumors display molecular, biological and clinical heterogeneity. Using bulk geneexpression profiling, DLBCL was stratified into two molecular subtypes based on their cell of origin (COO): germinal center B cells (GCB) or activated B cells (ABC)^7^. While this stratification identified patients with inferior outcomes after R-CHOP, it was not sufficient to predict the efficacy of newer treatment regimens^8–10^. Whole exome sequencing deciphered the genetic heterogeneity of DLBCL and classified tumors into 5-7 genetic subclasses based on their mutational landscape^11–13^. Several methods have been applied to unravel heterogeneity in the tumor microenvironment (TME), including deconvolution methods of bulk transcriptomic analysis that identified different types of lymphoma microenvironments (LME1-4)^14^ and lymphoma ecotypes (LE1-9)^15^. Single-cell or single-nuclei RNA sequencing analysis have further highlighted heterogeneity across DLBCL and provided novel insight into the interactions between malignant and tumor-infiltrating cells^16–19^. Furthermore, spatial proteomics using high-parameter imaging have investigated the spatial organization of tumor and immune cells in DLBCL^20–24^.

A common strategy for immune escape is the loss of the major histocompatibility complex (MHC) that prevents T-cell recognition and cytotoxicity of tumor cells. DLBCL has a high frequency of MHC loss. By traditional immunohistochemical analysis, 40-60% of DLBCLs have been classified as MHC class I (MHC-I) negative and 20-40% as MHC class II (MHC-II) negative^25,26^. Relapsed DLBCL tumors harbor increased number of mutations in genes involved in antigen presentation, including *HLA-A, HLA-B and B2M*^27^, and cases in which the tumor cells scored positive for pan-HLA class I membrane expression showed significantly higher T-cell frequencies than cases with moderate or negative expression.^26^ This suggests that loss of antigen presentation is a common mechanism of T-cell-escape in DLBCL. Here, we used spatial proteomics to explore how tumor cell expression of MHC-I and MHC-II impact the TME of DLBCL and integrated the spatial data with gene mutations and proteomic profiles. Our results identified distinct neighborhood compositions surrounding MHC-I^pos^ versus MHC-I^neg^ tumor cells, with implications for choice of immunotherapy.

## Methods

See supplemental Methods for detailed protocols.

### Patient material

The study cohort consisted of patients with high-risk *de novo* DLBCL included in two Nordic clinical trials^28,29^ (Supplementary Table S1). All samples were diagnostic biopsies, see Supplementary Fig. S1 for an overview. The study was approved by the regional committee for medical and health research ethics in Norway (ID 23192).

### Imaging Mass Cytometry analysis

Imaging mass cytometry (IMC) was performed as previously described^30^ on 2-μm tissue microarray (TMA) sections, stained with 31 antibodies (Supplementary Table S2), using the Hyperion Imaging System (Standard Biotools). Single-cell segmentation was done using IMCtools and customized CellProfiler modules from the Bodenmiller lab^31^, and was manually inspected for optimal segmentation. Following tSNE-CUDA and FlowSOM analyses (CellMass Cytobank), clusters were annotated by lineage markers: B cells/tumor (CD20, CD45RA, FOXP1), T cells (CD3, CD4, CD8, CD45RO, GzmB), macrophages (CD68, CD163, CD16), endothelium (CD34) and stroma (collagen) (Supplementary Fig. S2).

To identify different neighborhoods, the images were analyzed in CytoMAP^32^ using a raster scan with 20-μm-radius, followed by self-organizing map (SOM) clustering of the cell type abundances within in each neighborhood. The prevalence of neighborhood types in each sample was hierarchically clustered using R.

### Targeted DNA sequencing, proteomic profiling and available datasets

Targeted sequencing of formalin-fixed paraffin-embedded (FFPE) samples from 31 of the patients used an Agilent SureSelect custom design of 139 lymphoma relevant genes^27^. In addition, data from targeted sequencing of cell-free DNA was available from 21 of the patients, of which 13 patients had data from both FFPE and cell-free DNA^33^. Proteomic data from FFPE tumor tissue obtained by label-free quantification nano liquid chromatography-tandem MS (LFQ nLC-MS/MS) was available from 32 of the patients^34^. Serum protein profiling by Olink Explore 1536 library was available from 15 of the patients^35^. For validation, single-cell transcriptomes were available from Ye *et al*.^16^ MHC-I gene expression was inferred from bulk RNA cohorts (Chapuy *et al*.^12^, Schmitz et al.^13^) by application of CIBERSORTx^15^ in high-resolution mode with a B-cell–focused reference signature to infer sample-level immune-cell fractions. MHC class I (*HLA-A, HLA-B, HLA-C, HLA-E*) expression was extracted from imputed B-cell matrices.

## Results

### Contrasting tumor microenvironment types in DLBCL

To characterize the TME in DLBCL, we performed IMC analyses of diagnostic biopsies from patients included in the clinical trials NLG-LBC-04 and NLG-LBC-05^28,29^. These trials enrolled younger, previously untreated, high-risk patients diagnosed with large B-cell lymphoma for treatment with dose-densified chemo-immunotherapy combined with early systemic CNS prophylaxis. All patients with available FFPE tissue and a representative tumor area in TMAs were included for IMC. After optimal single-cell segmentation and quality control, more than 0.9 million single cells were clustered using FlowSOM and manually annotated, identifying eight cell types across the 67 patients which also included anaplastic and T-cell/histiocyte rich DLBCL, follicular lymphoma grade 3B and primary mediastinal B cell lymphoma (PMBCL) (Supplementary Fig. S2F). These cases with diagnoses other than DLBCL not otherwise specified (NOS) served as controls and were excluded from further downstream analyses, yielding a final DLBCL NOS cohort of 53 patients. Of these, 68% had an age-adjusted international prognostic index score of 2, 58% were GCB type COO and the median age was 54 years (Supplementary Table S1). The median tumor cell content was 76%, and CD8 T cells (4.5%) and M1 macrophages (3.1%) were the most common immune cell types (Fig. 1A). Patients with multiple TMA cores generally demonstrated low spatial heterogeneity (Supplementary Fig. S3A).

**Figure 1:**
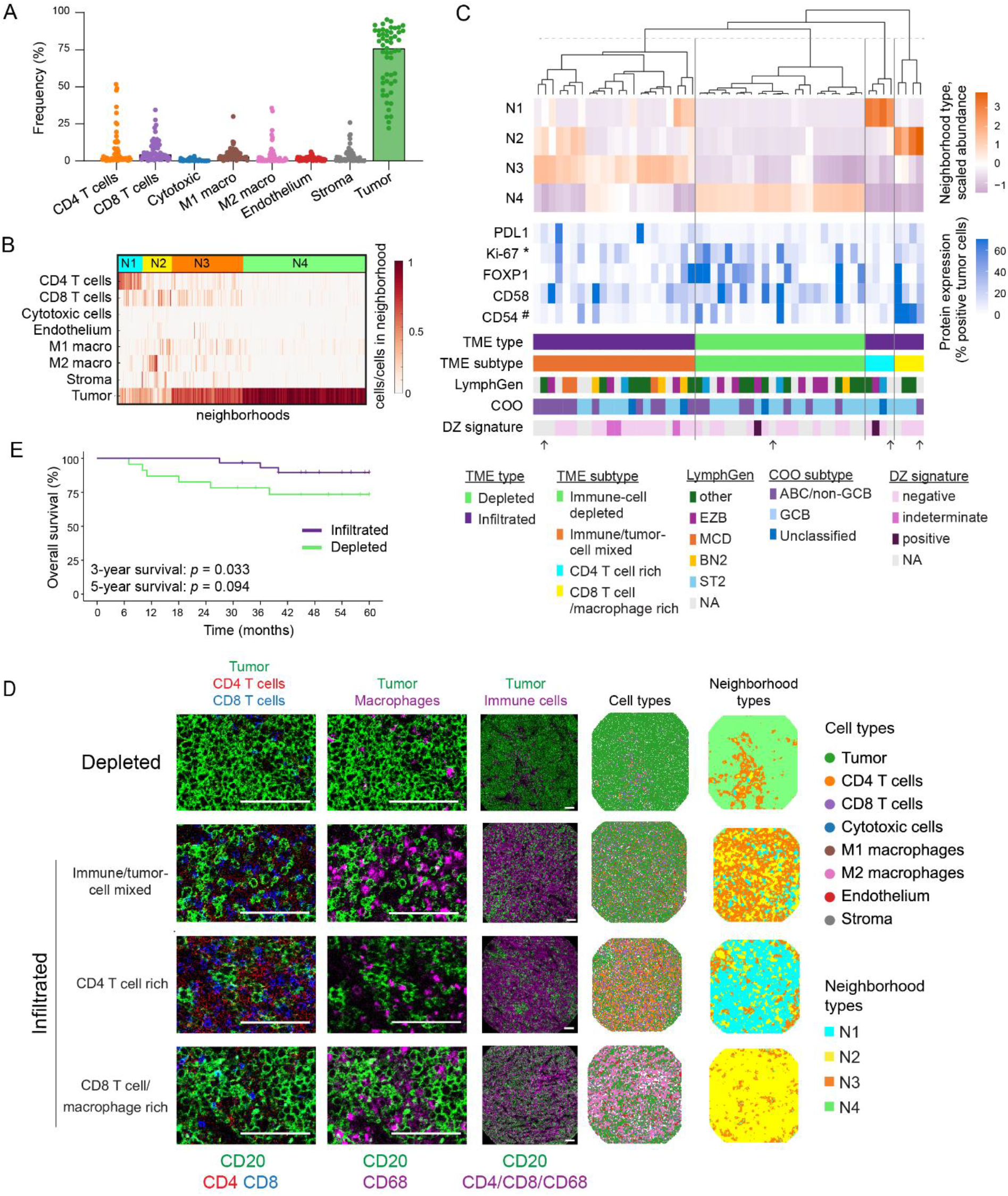
Identifying types of TME in DLBCL. 53 DLBCL NOS were analyzed by IMC. **A)** Frequency of cell types after segmentation and annotation. Bars indicate median. **B)** Raster-based 20-μm neighborhoods were SOM clustered based on cell-type composition using CytoMAP^32^. **C)** Hierarchical clustering of neighborhood prevalence in each sample identified four clusters, one immune-cell depleted and three immune-cell infiltrated, together annotated as infiltrated. Arrows indicate representation cases shown in panel D. Tumor-specific protein expression, in addition to cellof-origin and LymphGen classification, are shown for each case. * *p* < 0.01, Mann-Whitney test between the two TME types. # *p* < 0.01, Kruskal Wallis and Dunn’s multiple comparison test of the four TME subtypes. **D)** One example of each TME subtype shown as IMC pseudo images colored by marker expression, cell type maps colored per cell type as indicated, and neighborhood maps colored by neighborhood type from the SOM clustering in 1B. White scale bars indicate 100 μm. **E)** Kaplan-Meier plot of overall survival in patients with depleted and infiltrated DLBCL.

To identify differences in spatial immune cell composition, we performed a raster scan across all images, counting cell types present within 20-µm-radius neighborhoods. Clustering of neighborhoods revealed four types, N1-N4 with N1 and N2 containing the highest frequency of immune cells and consequently the lowest tumor cell content, followed by mixed cellular composition in N3 and N4 dominated by tumor cells (Fig. 1B). N1 neighborhoods had a large proportion of CD4 T cells, and N2 were rich in CD8 T cells and macrophages (Fig. 1B). Hierarchical clustering of the prevalence of neighborhood types identified four TME subtypes: an immune-cell-depleted subtype dominated by N4 (43.5%), immune- and tumorcell-mixed subtype dominated by N3 (41.5%), CD4-T-cell-rich TME subtype dominated by N1 (7.5%), and a CD8-T-cell/macrophage-rich subtype dominated by N2 (7.5%) (Fig. 1C, D; Supplementary Fig. S3B). The three TME subtypes with varying degrees of immune-cell infiltration were combined to give two main TME types, infiltrated and depleted. The TME subtypes showed no clear association with tumor COO classification or LymphGen genotypes (Fig. 1C, Supplementary Fig. S3C), and the TME types were not significantly associated with any recurrent somatic mutations (not shown). The frequency of tumor cells expressing the proliferation marker Ki-67 was higher in depleted cases (*p* = 0.04), whereas FOXP1, a transcription factor involved in early B-progenitor and germinal-center B-cell development,^36^ was more often expressed in depleted cases within the ABC molecular subtype (*p* = 0.03; Fig. 1C, Supplementary Fig. S4). CD54^pos^ tumor cells were significantly enriched in cases of the CD8-T-cell/macrophage-rich TME subtype (*p* = 0.02, Fig. 1C, Supplementary Fig. S4). The tumor-cell expression of immune checkpoints PD-L1 and CD58, a ligand for the T-cell coactivator CD2, was not associated with TME types (Fig. 1C). PD-L1 was more often expressed in macrophages than tumor cells (Supplementary Fig. S4), as previously described^37^. The depleted cases had significantly shorter overall survival compared to the infiltrated cases (Fig. 1E), in line with previous studies reporting poor survival in B-cell-enriched DLBCL^14,15^. In a univariate Cox proportional hazards model of each cell type, the frequency of M1 macrophages was borderline significant (*p* = 0.06), and cases with M1-macrophage frequency above the cohort median had better progression free survival (*p* = 0.024) (Supplementary Fig. S5).

### Proteomic signatures of TME types

High-throughput mass-spectrometry-based proteomic data obtained from FFPE bulk tissue analyses were available for 32 of the patients^34^. To further characterize depleted versus infiltrated DLBCL tumors, we performed a targeted re-evaluation of protein expression focused on the new classifications. Applying a threshold of at least 50% difference in expression, 157 proteins were significantly upregulated in infiltrated cases, and 122 proteins were significantly upregulated in depleted cases (Fig. 2A, *p* < 0.05). Concordant with higher expression of Ki-67 in depleted DLBCL (Fig. 1C, Supplementary Fig. S4), this TME type had higher expression of several proteins associated with proliferation including MCM5, FAM129A, and IGF2BP3 (Fig. 2A). MCM5 (minichromosome maintenance 5) showed the highest differential expression, and all six members of the MCM2-7 complex were among the 22 most upregulated proteins. The MCM2-7 complex is essential for DNA replication and is linked to increased proliferation in cancer^38^. Upregulation of *MCM5* in tumor-rich DLBCL was also seen in malignant B cells from a single-cell RNA sequencing (scRNA-seq) data set^16^, indicating that the enriched proteins in the bulk proteomic data are tumor-cell specific (Fig. 2B). Two proteins involved in MHC-II antigen presentation, CD74 and HLA-DQA, were also upregulated in depleted cases, and *CD74* gene expression was validated in the scRNA-seq data set (Fig. 2B). CD74 (invariant chain) is a chaperone needed in the assembly and trafficking of MHC-II. In data from the Tumor Cancer Genome Atlas program (TCGA), *CD74* mRNA is overexpressed in DLBCL compared to healthy tissue, and CD74 copy number variations have a prognostic significance in DLBCL^39^. Consistent with our IMC data, the monocyte/macrophage-associated proteins CD163 and CD14 were increased in infiltrated cases. Interestingly, hierarchical clustering of the significant differentially expressed proteins revealed two main clusters, of which one cluster contained 11 out of 12 cases classified as depleted, highlighting the biological difference between these two groups (Fig. 2C).

**Figure 2:**
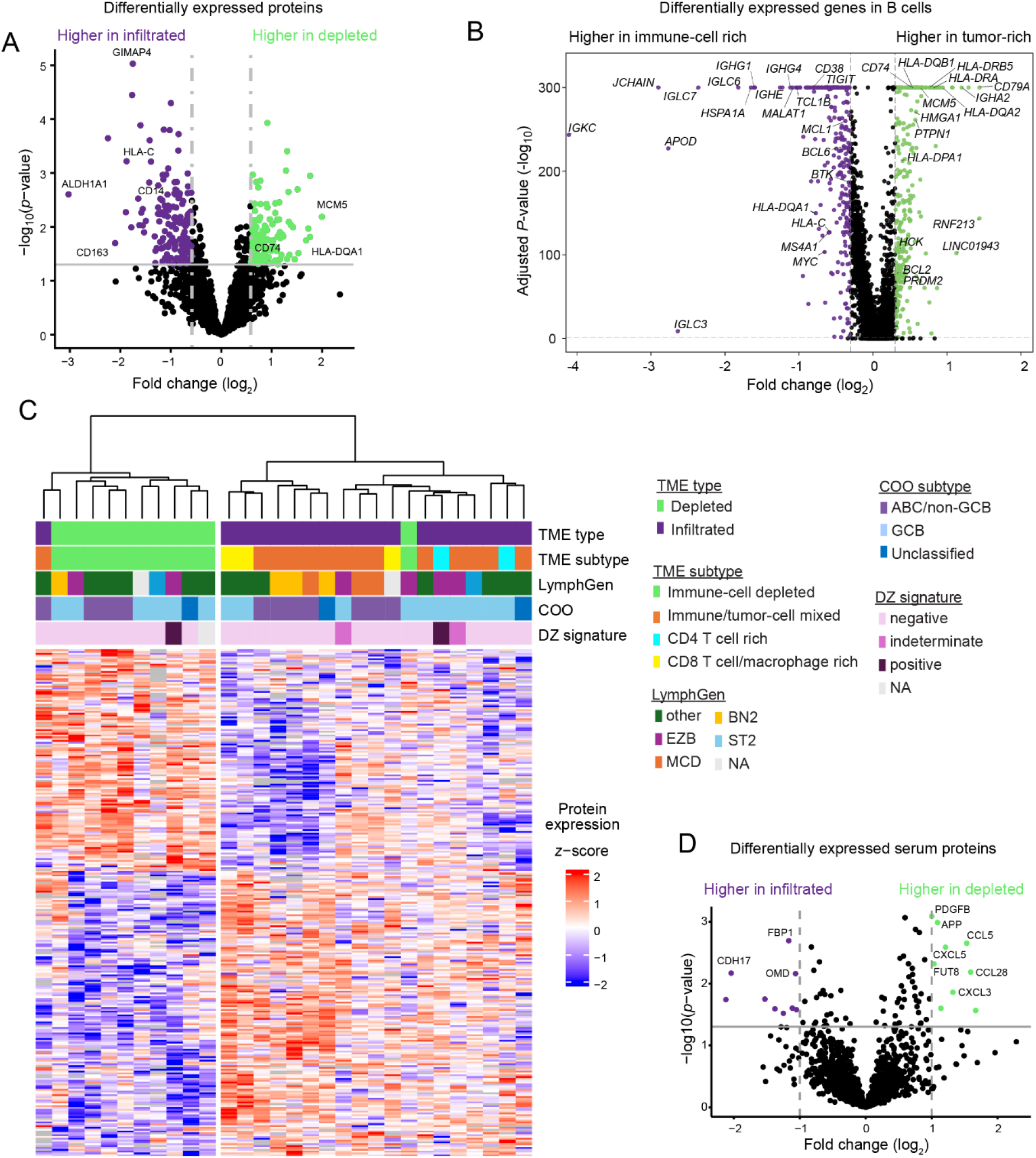
Differentially expressed proteins between TME groups. **A)** FFPE samples from 32 of the 53 patients in Figure 1 were analyzed by mass spectrometrybased proteomics^34^. Volcano plot showing differentially expressed proteins between depleted and infiltrated DLBCL (vertical lines indicate log_2_ fold change of 0.585 (minimum 50% difference in expression), horizontal line indicates *p* = 0.05). **B)** Differentially expressed genes from single-cell RNA sequencing data set^16^. Tumor-cell enriched cases were defined as having more than 75% B cells (Supplementary Figure 6B). Vertical lines indicate log_2_ fold change of 0.3, horizontal line indicates *p* = 0.05. **C)** Dendrogram of hierarchical clustering with input of the 279 differentially expressed proteins in (A). **D)** Serum samples from 15 of the 53 patients were analyzed by Olink multiplexed protein analysis. The volcano plot shows differentially expressed proteins in depleted vs. infiltrated cases (vertical lines indicate log_2_ fold change of 1, horizontal lines indicate *p* = 0.05).

Pathway enrichment analysis of proteomic data showed enrichment of cell cycle and DNA replication pathways in depleted DLBCL (Supplementary Fig. 6A). Gene set enrichment analysis of malignant B-cells from the scRNA-seq data showed upregulation of nucleic acid biosynthesis pathways in tumor-rich cases and enrichment of immune response pathways in cases with higher immune cell infiltration, indicating interaction between lymphoma cells and other immune cells of the TME (Supplementary Fig. 6C).

To connect the TME proteomics with blood-based clinical assays, serum proteome profiling was performed for the NLG-LBC-05 trial patients.^35^ Fifteen of these overlapped with the IMC cohort. Analysis of differentially expressed serum proteins revealed that several chemokine ligands including CXCL3, CXCL5, CCL5 and CCL28 were upregulated in depleted compared to infiltrated cases (Fig. 2D). A cluster analysis in the original publication identified a subset of patients with a serum profile characterized by lymphocyte activation (cytolytic proteins, checkpoint molecules) and inflammation (IL-10, IL-18, IFNγ),^35^ consistent with a T-cell response in the TME. These patients were associated with increased T-cell exhaustion in the TME and poor outcome^35^. Three of the overlapping patients belonged to the inflamed cluster, and these were among the infiltrated cases (Supplementary Fig. S3A).

### Common loss of MHC-I in depleted cases

Since loss of MHC-I and MHC-II are common immune-escape mechanisms in DLBCL^25,26^, we investigated expression of HLA-A and HLA-B (denoted MHC-I) and pan-MHC-II in relation to TME subtypes. We observed large heterogeneity in tumor-cell MHC-I expression across cases, and most cases contained tumor cells with MHC-I loss (Fig. 3A and B). Based on Youden index analysis to identify the optimal cut-point relative to TME type, cases with >52.05% of the tumor cells expressing MHC-I were defined as MHC-I positive, and those with ≤52.05% as MHC-I negative. Interestingly, MHC-I negative tumors were enriched for the depleted TME type (Fig. 3A, *p* = 0.04) and had significantly fewer CD4 and CD8 T cells but not M1 or M2 macrophages (Supplementary Fig. S7A). In contrast, there was no optimal cutpoint for tumor-cell expression of MHC-II that were associated with TME type (Supplementary Fig. S7B). Our cohort lacked statistical power to identify patterns between TME subtypes and gene mutations associated with loss of MHC-I, but we noted that two cases with almost complete loss of MHC-I had *B2M* mutations (Fig. 3A). MHC-I gene mutations could not be assessed due to lack of matched germline samples. Most MHC-I negative cases also had low frequency of MHC-II^pos^ tumor cells (Fig. 3B).

**Figure 3:**
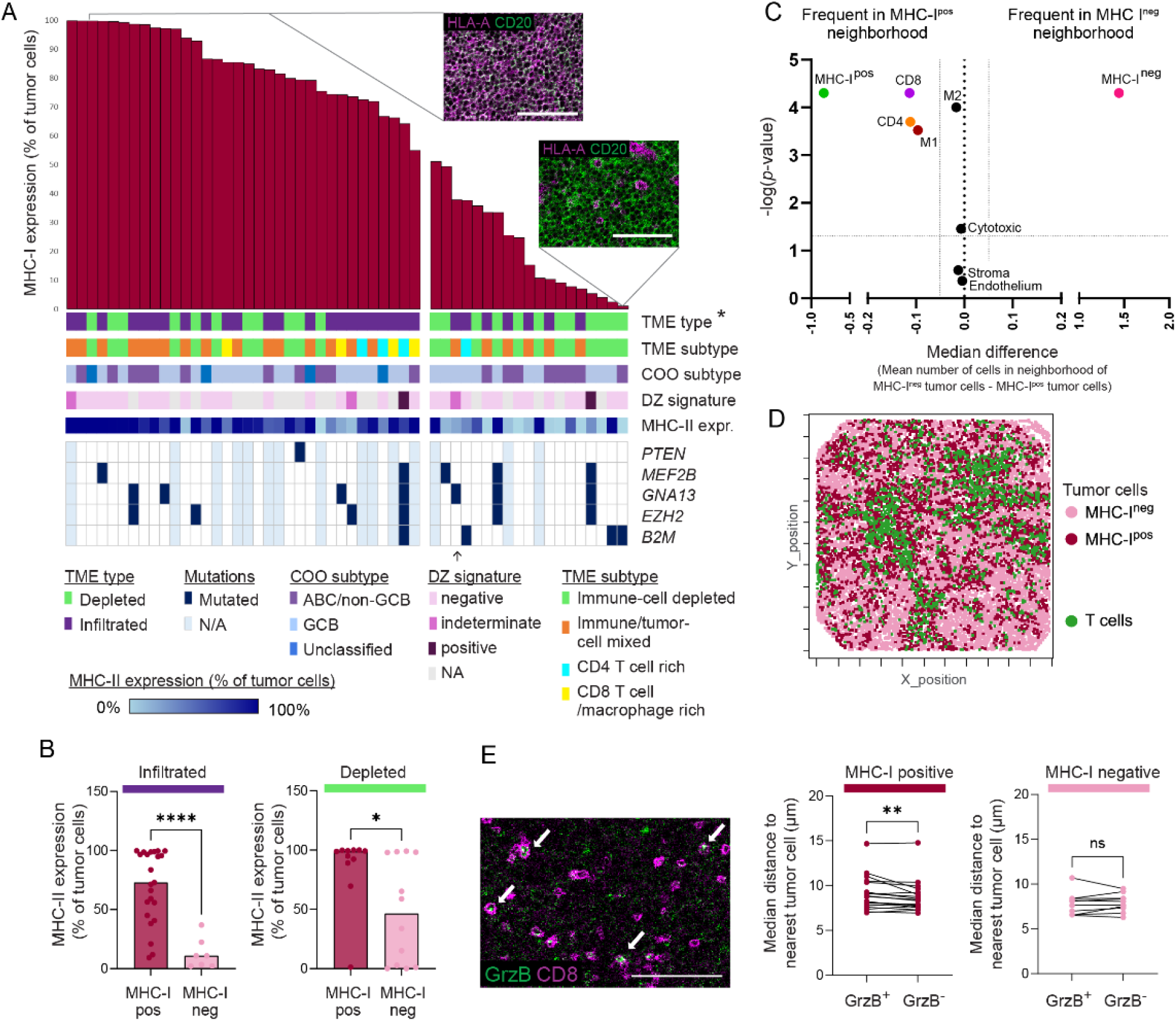
MHC class I expression is associated with TME type. **A)** HLA-A and HLA-B expression (denoted MHC class I (MHC-I) expression) was measured by IMC. All samples were ordered by frequency of tumor cells expressing MHC-I. Youden index was used to find optimal cut-point for loss of MHC-I expression relative to TME type. Bars below the graph indicate TME type, TME subtype, COO subtype and mutational status of genes previously shown to be associated with loss of MHC-I. * *p* < 0.05 in Fisher exact test. Inserts show example images of tumor cells with expression (top) or loss (bottom) of MHC-I. **B)** MHC-II expression plotted in MHC-I negative and positive cases separately for both depleted and infiltrated cases. * *p* < 0.05; **** *p* < 0.0001 in Mann-Whitney test. **C)** In cases with heterogeneous MHC-I expression (20-80% of tumor cells were MHC-I^pos^), the cells within a radius of 20 μm from each tumor cell were counted. For each cell type, the difference of mean number of cells in the neighborhoods of MHC-I^neg^ and MHC-I^pos^ tumor cells was calculated. The median difference for each sample and the *p*-value from a Wilcoxon matched-pairs signed rank test were plotted in a volcano plot. **D)** Spatial organization of MHC-I^pos^ and MHC-I^neg^ tumor cells relative to T cells in one infiltrated case. **E)** Image shows CD8 T cells expressing Granzyme B. White scale bars indicate 100 μm. The distance from Granzyme B^+^ and Granzyme B^-^ CD8 T cells to the nearest tumor cell is calculated and plotted separately for MHC-I positive and negative cases. ** *p* < 0.01 in Wilcoxon test.

To investigate if the depletion of immune cells was also seen in the neighborhoods of MHCI^neg^ tumor cells, we performed further spatial analysis of cases with heterogeneous MHC expression (20-80% MHC-I^pos^ tumor cells). The neighborhoods of MHC-I^pos^ tumor cell were enriched for other MHC-I^pos^ tumor cells, CD4 and CD8 T cells and M1 macrophages (Fig. 3C). In contrast, MHC-I^neg^ tumor cells were typically found close to other MHC-I^neg^ tumor cells (Fig. 3C). One representative case illustrates that T cells are localized closer to MHC-I^pos^ tumor cells than to MHC-I^neg^ tumor cells (Fig. 3D, Supplementary Fig. S7C).

Further, we hypothesized that activated, Granzyme B^pos^ CD8 T cells are closer to tumor cells than Granzyme B^neg^ CD8 T cells when the tumor cells express MHC-I. Contrary to this, in MHC-I-positive cases the median distance from Granzyme B^pos^ CD8 T cells to the nearest tumor cell was significantly larger than from Granzyme B^neg^ CD8 T cells to the nearest tumor cell, although the differences were small (Fig. 3E). In MHC-I negative cases, we found no significant difference between Granzyme B^pos^ and Granzyme B^neg^ CD8 T cells (Fig. 3E).

To validate the association between tumor-cell MHC-I loss and reduced immune-cell infiltration, we applied CIBERSORTx to annotate and enumerate cell-type frequencies and to impute B-cell transcriptomes in two publicly available bulk RNA-sequencing datasets^12,13^. In each data set, we stratified the cases by B-cell fraction and compared the MHC-I gene expression between cases with the 20% highest and 20% lowest frequency of B cells (Supplementary Fig. S8). *HLA-B* expression was significantly lower in B cells from cases with the highest B-cell content (top 20%) in both data sets, hence supporting our results from the IMC analysis that loss of MHC-I is one of the major underlying mechanisms limiting T-cell infiltration.

## Discussion

In this study, we provide insight into how tumor-cell MHC-I status impacts the spatial architecture of *de novo* DLBCL tumors. We identified four distinct TME subtypes: one immune-cell depleted and three subtypes of infiltrated TME, similar with previous studies identifying both immune-cell-rich and immune-cell-sparse DLBCL^20,22^. The TME subtypes were observed across DLBCL genetic classes and mutational profiles. No robust association between DLBCL TME subtypes and genetic subtypes has so far been found^16–19^. Our data points to tumor MHC-I status as a main factor driving immune infiltration. DLBCL cases categorized as MHC-I negative were associated with the depleted TME type, which is concordant with prior studies showing that loss of MHC-I was associated with lower T-cell infiltration^23,25,26^. In cases with heterogeneous MHC-I expression, we further demonstrated that the neighborhoods of MHC-I^pos^ tumor cells were enriched for T cells and M1 macrophages, contrasting the neighborhoods of MHC-I^neg^ tumor cells which were immunecell depleted. Downregulation or loss of MHC-I is a common mechanism of resistance to immune checkpoint blockade in cancer^40–42^, and likely a major underlying cause for limited clinical benefit to this immunotherapeutic approach in DLBCL.

Downregulation or loss of MHC-I in DLBCL has been linked to recurrent mutations in the antigen presentation machinery, including *B2M*, required for the assembly and stability of MHC-I, and loss of heterozygosity and mutations in the *HLA* genes themelves^12,13,25,27^. Such mutations seem to be a major cause for loss of MHC-I in DLBCL, as somatic inactivation of *B2M* and the *HLA-I* loci were found in 80% of DLBCL cases that were scored as MHC-I negative as estimated by lack of membrane staining in >90% of tumor cells^43^. In addition to somatic mutations, downregulation of MHC can be caused by epigenetic silencing. Lymphoma cells harboring the *EZH2* Y641 mutation had reduced expression of MHC-I and MHC-II as compared to *EZH2* WT cells^25^. Downregulation was linked to repression of the MHC transactivators *NLRC5* and *CIITA* and could be reversed by use of EZH2 inhibitors^25^. EZH2 as part of the polycomb repressor complex can silence multiple genes, and EZH2 inhibitors were shown to sensitize tumor cells to CAR T-cell killing in hematological and solid tumor models by upregulating genes involved in T-cell co-stimulation and IFN-γ-regulated genes in addition to *HLA* genes^44^. Further evidence for a role of tumor-cell MHC status in determining the efficacy of CAR T-cell therapy comes from a recent study, in which endogenous bi-allelic loss of MHC-I and CD58 in tumor cells were associated with resistance to CD19 CAR T-cell therapy^45^.

The relevance of TME-typing for immunotherapy including CAR T-cell treatment was recently demonstrated by Li *et al*.^19^ By focusing on non-tumor TME cells, they described three DLBCL microenvironment archetype profiles: A lymph node (LN) archetype that is T-cell supporting and most closely aligned with normal LN architecture, an inflamed archetypes where the T cells were suppressed (TEX), and a T-cell excluding archetype with high frequencies of cancer-associated fibroblasts and tumor-associated macrophages (FMAC) ^19^. Interestingly, DLBCL patients with the LN archetype demonstrated highest CAR T-cell efficacy as compared to standard of care, which contrasted the TEX archetype with no benefit of CAR T-cell therapy ^19^. As tumor cells were not part of the archetypes, the relationship to tumor cell abundance is not clear, nor the tumor-cell expression of MHC-I.

Our study analyzed diagnostic biopsies from a cohort of uniformly treated patients. However, a limitation is the small patient cohort, reducing statistical power when analyzing patient level factors. Furthermore, the IMC analysis did not have sufficient resolution to reliably distinguish between intracellular and membrane staining of MHCs, and the fraction of MHC-I cell surface negative tumor cells might therefore be underestimated. Despite these limitations, we have identified an association between immune-cell infiltration and MHC-I expression. In the future, TME-based classification of tumor specimens that incorporates tumor-cell MHC-I status may improve the selection of immunotherapies.

