## Supplemental information for "Immune-cell depleted diffuse large B-cell lymphomas have reduced expression of MHC class I"

### Supplementary methods

#### Patient material

We included patients with high-risk de novo DLBCL from two Nordic phase II clinical trials (CRY-04/NLG-LBC-04/NCT01502982[1] and CHIC/NLG-LBC-05/NCT01325194[2]). Patients were below 65 years with an age-adjusted International Prognostic Index (IPI) score of 2-3 and/or increased risk of central nervous system (CNS) recurrence. All received biweekly R-CHOP with etoposide and systemic CNS prophylaxis.

Tissue microarrays (TMAs) of diagnostic formalin-fixed paraffin-embedded (FFPE) biopsies were made. All TMA cores were reviewed by a pathologist, ensuring that the imaged cores were DLBCL not otherwise specified (NOS) (WHO classification, 2008[3]) and representative of the tumor area. Cell of origin (COO) was determined by the DLBCL90 assay[4] in 33 of the patients and by Hans' algorithm for the remaining 20.

#### Imaging Mass Cytometry analysis

##### *Staining and acquisition*

Imaging mass cytometry (IMC) was performed as previously described[5] on 2-µm FFPE tissue section of the TMAs. Antigen retrieval was performed at high pH (Basic Antigen Retrieval Solution, R&D systems), and staining was performed in two steps with CD56-staining in room temperature for 5 hours followed by staining with a cocktail of all remaining antibodies (supplemental Table S2) at 4°C overnight. The following day, the slide was incubated with Cell-ID Intercalator-Ir (0.3 µM). The sections were ablated using the Hyperion Imaging System (Standard Biotools).

All antibodies were validated prior to staining, but some antibodies did not consistently stain the tissue and were not used for any biological interpretation (CD56, FOXP3, OX40, LAG3, TIGIT, PD1, ICOS).

##### *Cell segmentation*

The following segmentation pipeline was inspired by the Bodenmiller pipeline and used IMCtools and customized Bodenmiller CellProfiler modules[6]. All images were evaluated separately, and settings were adjusted for optimal segmentation.

MCD files were converted to ome.tiff files and two image stacks were made – one with all markers and one analysis stack with selected markers (Nuclear: Ir191, HistoneH2; Cytplasmic/membrane: CD20, CD45, CD45RO, CD45RA) for pixel classification in Ilastik. In

CellProfiler 3.1.8, the image stacks were filtered using the Smooth Multichannel module to remove outlier pixels. Random crops were made to save computing power during training in Ilastik.

Pixel classification of the analysis stack was performed in Ilastik 1.3.0. Three classes of pixels were defined: nucleus, cytoplasm/membrane and background. Training was primarily performed on the cropped analysis stacks and then applied to the original analysis stack by batch processing. The probability maps were evaluated separately and if needed, training was re-done on the full analysis stack.

Segmentation was performed in CellProfiler. The probability map stacks were split in nuclear, cytoplasm/membrane and background. Nuclei were defined by the module "Identify primary object" and the Otsu thresholding method (threshold correction factor was adjusted separately). Objects outside the typical diameter (3-20 pixels) were excluded. Distinguishing between clumped objects was performed with the Laplacian of Gaussian and propagation method, except a few images where the shape method gave better results (distance for suppressed local maxima was adjusted). Cells were identified by expanding 2-4 pixels from the nucleus and the probability map of cytoplasm/membrane was used to visually determine the optimal number of pixels to expand.

Macrophages were segmented separately based on CD68 staining only. The raw CD68 image was fed into CellProfiler and multiplied by 1000-5000 depending on staining strength. Macrophages were defined by the module "Identify primary object" and the Otsu thresholding method (threshold correction factor was adjusted separately). Objects outside the typical diameter (5-50 pixels) were excluded. Cells with more than 50% overlap with a macrophage were excluded and remaining overlapping pixels were defined as macrophage before the two object masks were combined.

Image masks and single marker TIFF files for each image were opened in HistoCAT[7]. The quality of the mask was evaluated, and segmentation pipeline was adjusted if the mask was not satisfactory. Single cell events were exported as CSV files.

*Cell annotation*

tSNE-CUDA and FlowSOM analyses (CellMass Cytobank) were performed for clustering and clusters were annotated by lineage markers: B cells/tumor (CD20, CD45RA, FOXP1), T cells (CD3, CD4, CD8, CD45RO, GzmB), macrophages (CD68, CD163, CD16), endothelium (CD34) and stroma (collagen). Overclustering was followed by manually merging into the final cell types. Due to variation in signal intensity across images, the samples were divided into three intensity groups based on the combined 99$^{th}$ percentile intensity of CD45RO (expressed by most T cells) and CD68 (expressed by macrophages). The expression pattern in the cell types identified was similar between the intensity groups, but with an overall different scale (Suppl. Fig. 1). Tumor-cell specific protein expression was gated in Cytobank.

*Spatial analysis*

To identify different neigborhoods, single-cell tabulated data from the images were analyzed in CytoMAP[8] using a raster scan with 20-um-radius, followed by SOM clustering of the cell type abundances within in each neighborhood. The prevalence of neighborhood types in each sample was hierarchically clustered using R, revealing 4 TME subtypes.

Analysis of neighborhoods surrounding MHC $I^+$ and MHC $I^-$ tumor cells was performed in R and GraphPad Prism. For each image, the cells within 20 um of each tumor cell were counted and the mean number of each cell type surrounding MHC I+ and MHC I- tumor cells was calculated. A Wilcoxon matched-pairs signed rank test was performed for each cell type, pairing the mean number cells neighboring MHC I+ vs. MHC I- tumor cells. A volcano plot was made, showing p-value from the Wilcoxon test and the median difference of neighboring cells across images.

Analysis of distance from CD8 T cells to tumor cells was performed in R and plotted in GraphPad Prism. The FlowSOM clustering identified CD8 T cells and Granzyme $B^+$ cells as separate clusters. For this spatial analysis, the $CD8^+$ cells from the cluster of Granzyme $B^+$ cells were merged with the CD8 T cell cluster before Granzyme $B^-$ and Granzyme $B^+$ CD8 T cells were manually gated in Cytobank. The distance from each Granzyme $B^-$ and Granzyme $B^+$ CD8 T cell to the nearest tumor cells was calculated in cases classified as MHC class I positive and MHC class I negative.

**Targeted DNA sequencing**

Targeted sequencing used an Agilent SureSelect custom design of 139 lymphoma relevant genes.[9] To identify mutations, data from targeted sequencing and ctDNA sequencing (Meriranta *et al*.[10]) were combined. From ctDNA data, exonic and splicing aberrations were included, but non-synonymous mutations were excluded. A gene is considered mutated if it was scored as mutated in either of the two data sets.

**Lymphgen**

Mutations from targeted sequencing were identified with Mutect version 1. BCL6 and BCL2 transolcation status were determined from routine diagnostics. Copy number variations were identified from targeted sequencing using CNVkit fused lasso method. Focal aberrations were identified as any over 30 megabases while arm level copy number events were calculated as those with lengths greater than or equal to 70% of the cytoband chromosome length. For the ctDNA sequencing samples mutations were identified as previously described[10], no copy number status was included. Lymphgen status was determined from the online tool https://llmpp.ccr.cancer.gov/lymphgen/index.php version 2.0.

**Proteomic analyses**

Protein expression from FFPE tumor tissue obtained by label-free quantification nano liquid chromatography-tandem MS (LFQ nLC-MS/MS) was available from 32 patients. These data have been previously published[11]. Proteins differentially expressed between depleted and infiltrated cases were identified, and *p*-values were calculated by a two-tailed *t*-test. Proteins with an absolute $\log_2$ fold change greater than 0.585 and $p < 0.05$ have been highlighted. The KEGG PATHWAY database was used for pathway enrichment analysis. Proteins were ranked by a significance score calculated as sign($\log_2$ fold change) x -log10(*p* value) after filtering for absolute $\log_2$ fold change ≥ 0.3.

Serum protein profiling by Olink Explore 1536 library was available from 15 patients[12]. Differentially expressed proteins between depleted and infiltrated cases were calculated as difference between Normalized Protein Expression (NPX) values and *p*-values were calculated by a two-tailed *t*-test with a Benjamini-Hochberg correction for multiple testing with FDR of 0.05.

**Gene expression**

Single-cell RNA sequencing data (Ye *et al.*[13]) were used to identify differentially expressed genes between B-cell rich (defined as >75% B cells; $n = 3$) and immune-cell rich DLBCL cases ($n = 14$). The Gene Onotology Biological processes database was used for pathway enrichment analysis. Genes were ranked by a significance score calculated as sign($\log_2$ fold change) x -log10(*p* value) after filtering for absolute $\log_2$ fold change ≥ 0.3.

MHC class I gene expression was inferred from bulk RNA cohorts (Chapuy *et al.*[14], Schmitz *et al.*[15]). For each cohort, we applied of CIBERSORTx[16] in high-resolution mode with a B-cell–focused reference signature to infer sample-level immune cell fractions and B-cell–specific expression profiles. Within each cohort, samples were ranked by inferred B-cell fraction, and tumors in the top and bottom 20% quantiles were selected for comparison.

**Survival analyses**

The Kaplan-Meier method with log-rank test was used to estimate survival rates between different patient groups. Cox regression was used to estimate survival in univariable analyses.

#### Supplemental Figures

A

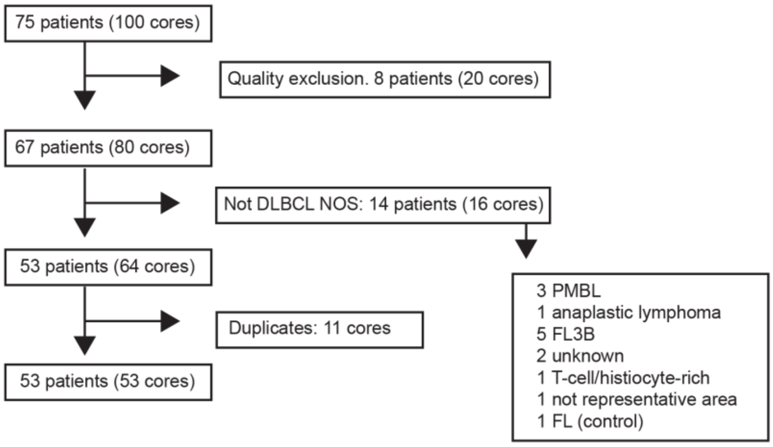


B

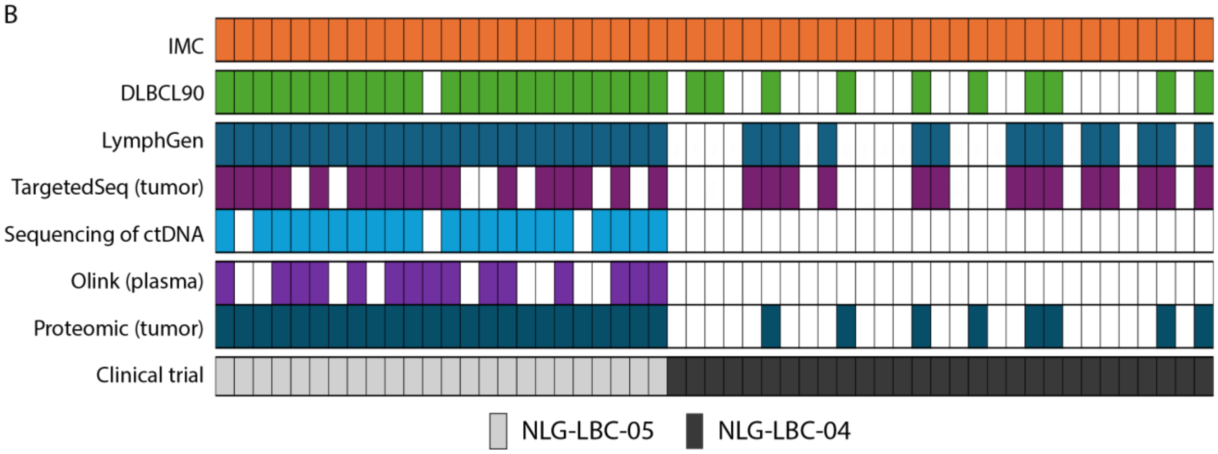


**Supplementary Figure S1.** Overview of samples

A) IMC analysis was performed on 99 TMA cores and one control sample, representing 75 lymphoma patients. 20 of these cores were excluded due to poor image quality. 16 cores were from patients with other diagnoses than DLBCL NOS as indicated. Of the remaining 53 patients, 6 of them had more than one core imaged. B) Overview of analysis types performed on samples from the 53 DLBCL NOS included in our IMC study.

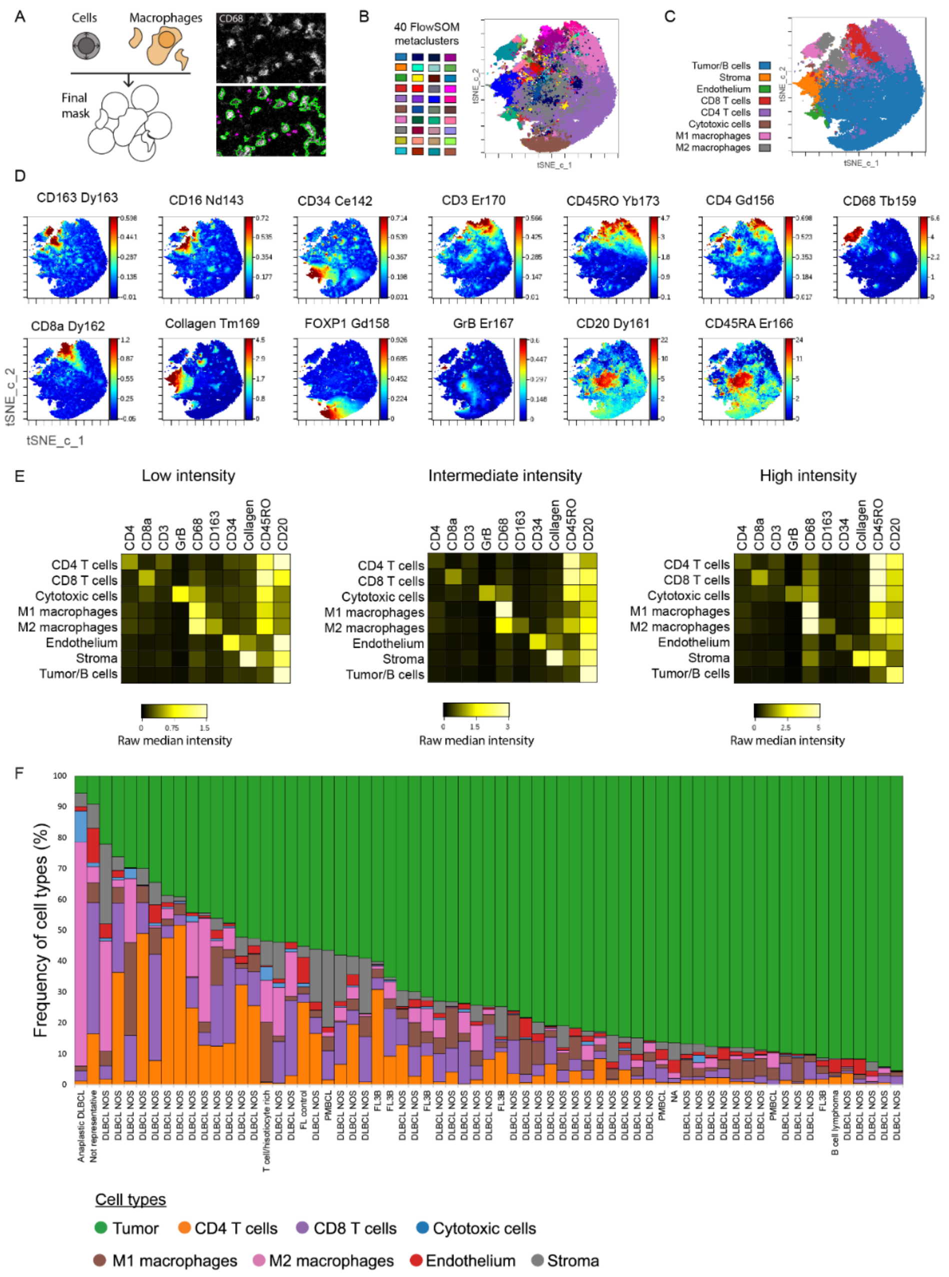

**Supplementary Fig. S2: Segmentation pipeline, clustering and annotation of cell types**

A) Segmentation pipeline. Nuclei were identified by nuclear markers (Ir191 and Histone H3 combined) and cells were identified by expanding 2-4 pixels from the nuclei. Macrophages were identified by CD68 expression independent of nucleus as these cells are large and the 2-µm section might not capture the nucleus. The cells and macrophage objects were combined for the final segmentation mask. The cells were overclustered by FlowSOM (B) before metaclusters were manually combined and annotated (C) based on marker expression (D). E) The samples were divided into intensity groups prior to FlowSOM to handle difference in intensity between samples. The same populations were identified in each group and had similar expression patterns, but with different relative intensity.  F) Samples ordered by percentage of tumor cells. DLBCL NOS compared to other types of lymphoma.

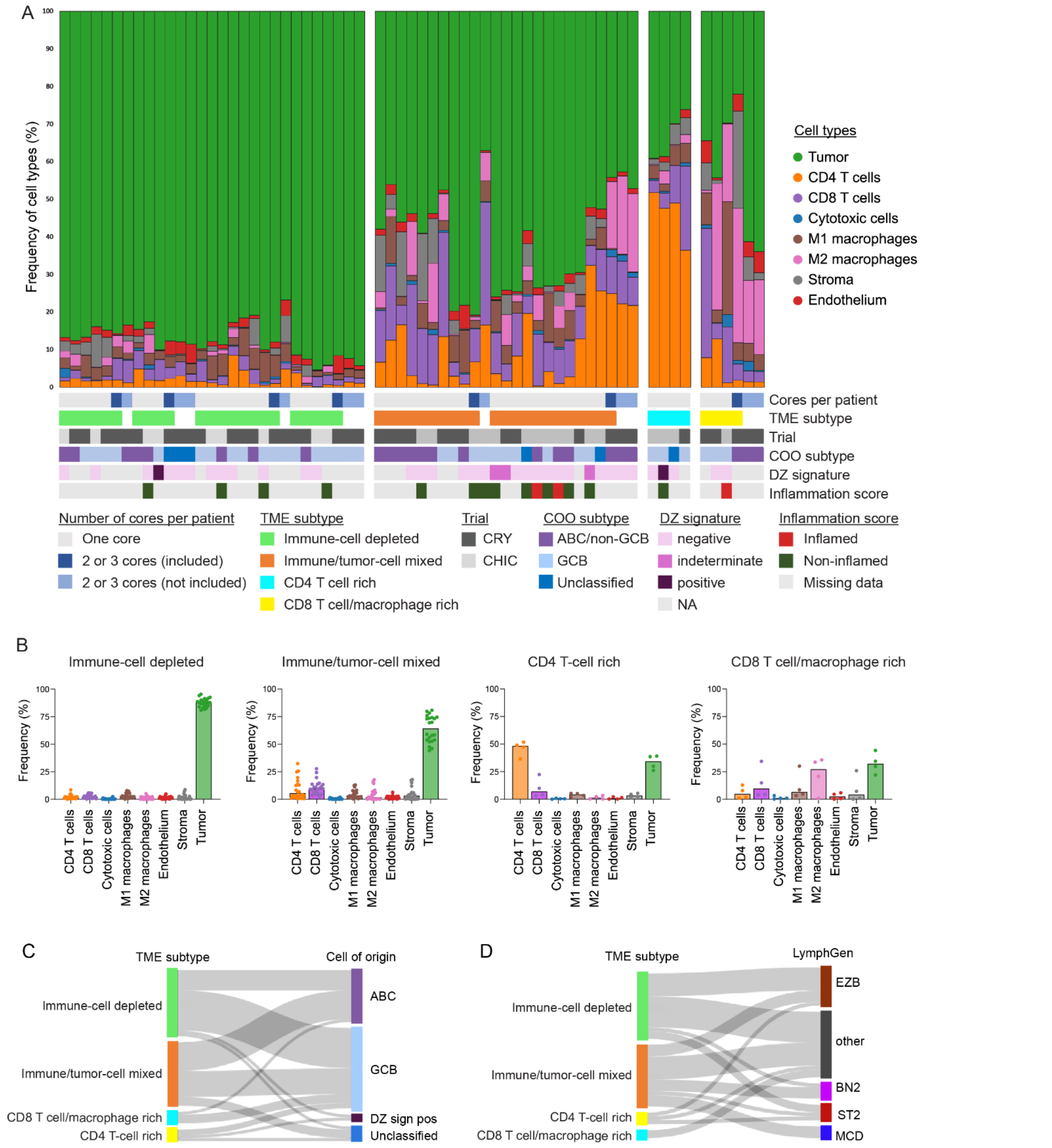


**Supplementary Fig. S3: Cell type frequency per sample in TME subtypes and spatial heterogeneity**

A) The 53 samples were ordered based on TME subtype. Seven of the samples had additional cores imaged and these are plotted next to the included cores (indicated in shades of blue). Bars also indicated COO subtype and inflammation score from Arffman *et al.*[12] B) Frequency of cell types separated based on TME subtypes. Bars indicate median. C) Sankey plot showing TME subtypes vs. cell-of origin. Cell-of-origin was determined by the DLBCL90-assay when available (*n* = 33) or by IHC analysis using the Hans' algorithm. D) Sankey plot showing TME subtype vs. LymphGen. LymphGen categories were determined based on the data from targeted or ctDNA sequencing[10].

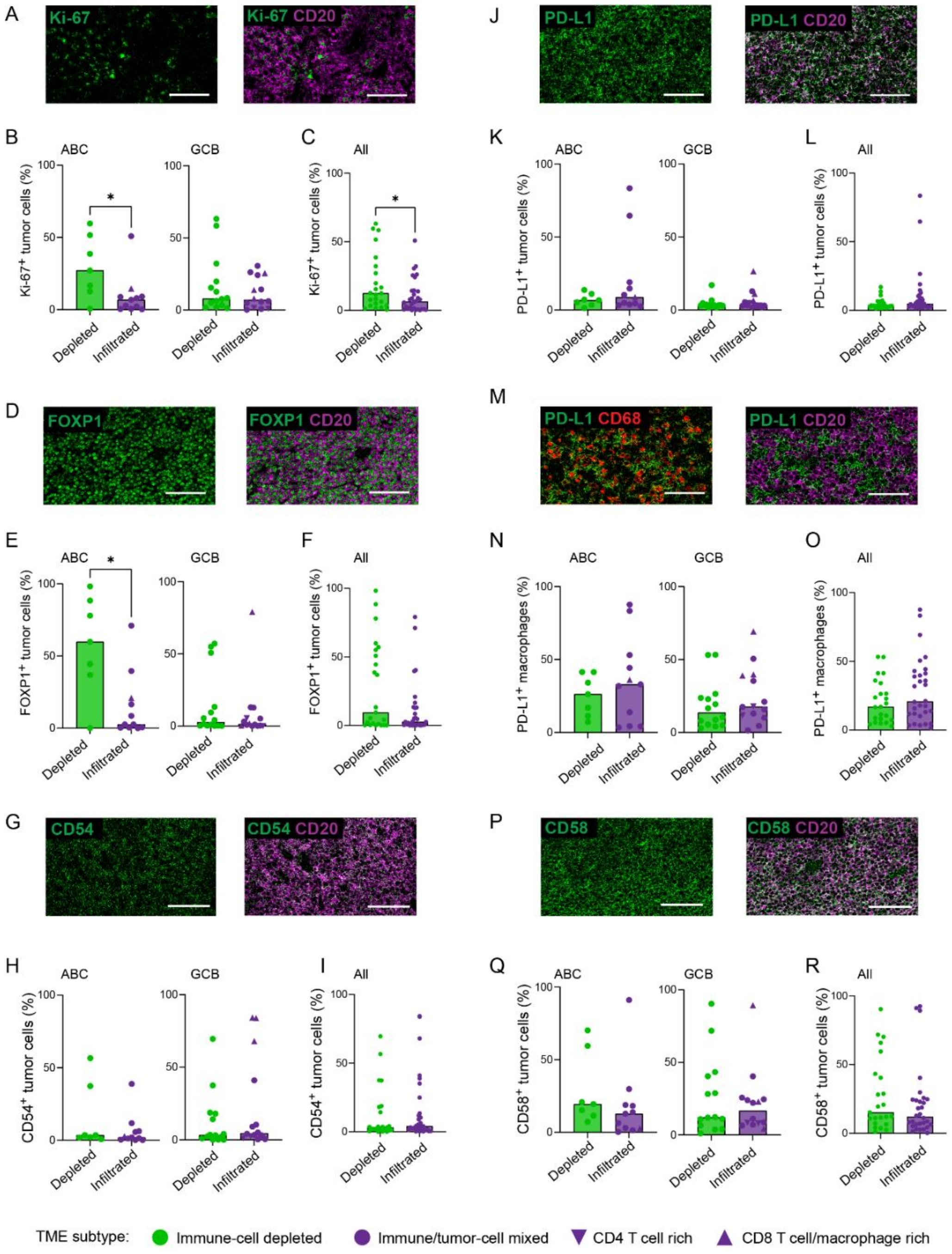


**Supplementary Figure S4: Tumor cell-specific protein expression differs between infiltrated and depleted DLBCL**

Example of Ki-67 (A), FOXP1 (D), CD54 (G), PD-L1 (J), CD58 (P) combined with CD20 expression in IMC images. Protein expression was determined by 2D-gating of positive cells. Frequency of tumor cells of ABC or GCB subtype expressing each protein (B, E, H, K and Q), and of all tumor cells expressing each protein in depleted and infiltrated cases (C, F, I, L, R). Example of a case where PD-L1 is primarily expressed by $CD68^+$ macrophages, not $CD20^+$ tumor cells (M), and frequency of PD-L1-expressing macrophages in cases of ABC or GCB (N) and in all cases (O). White scale bars indicate 100 µm. * indicates $p < 0.05$ in Mann-Whitney test.

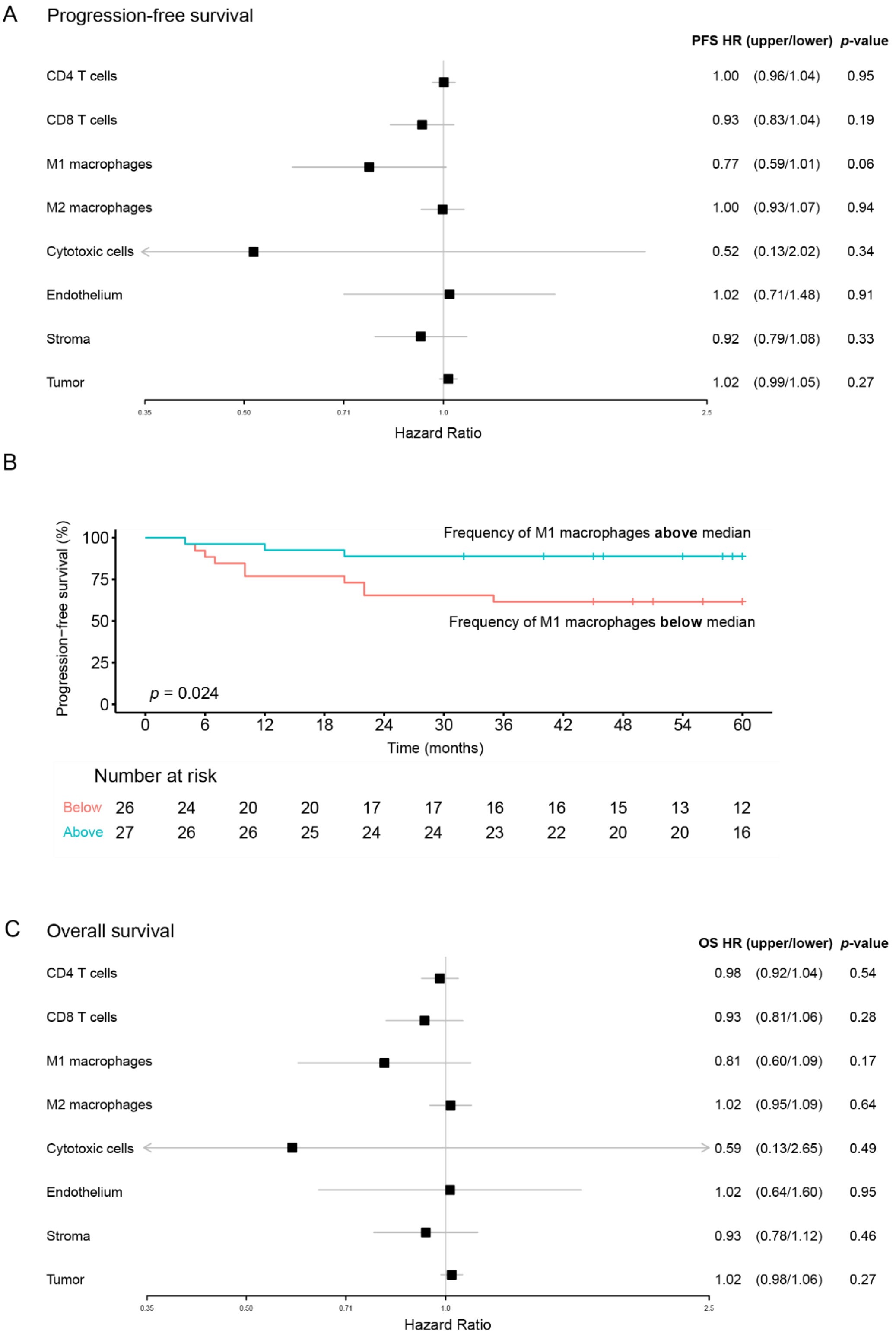


**Supplementary Fig. S5: Univariate Cox proportional hazards model**

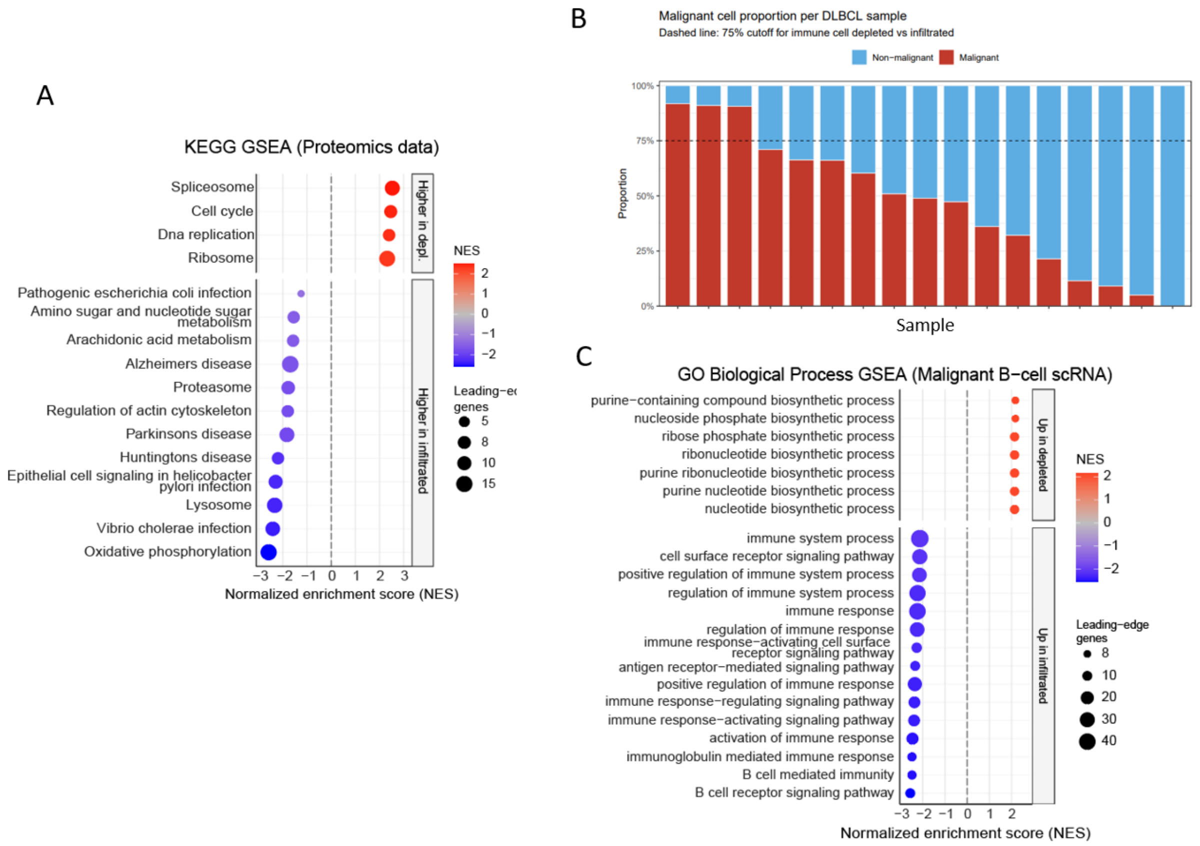


**Supplementary Fig. S6: Pathway enrichment analysis**

A) Pathway enrichment analysis of differentially expressed proteins from Figure 2A, using the KEGG PATHWAY database. B) Fraction of lymphoma cells in scRNA-seq data from *Ye et al*.[13]. C) Gene set enrichment analysis of differentially expressed genes from Figure 2B, using the GO Biological Process database.

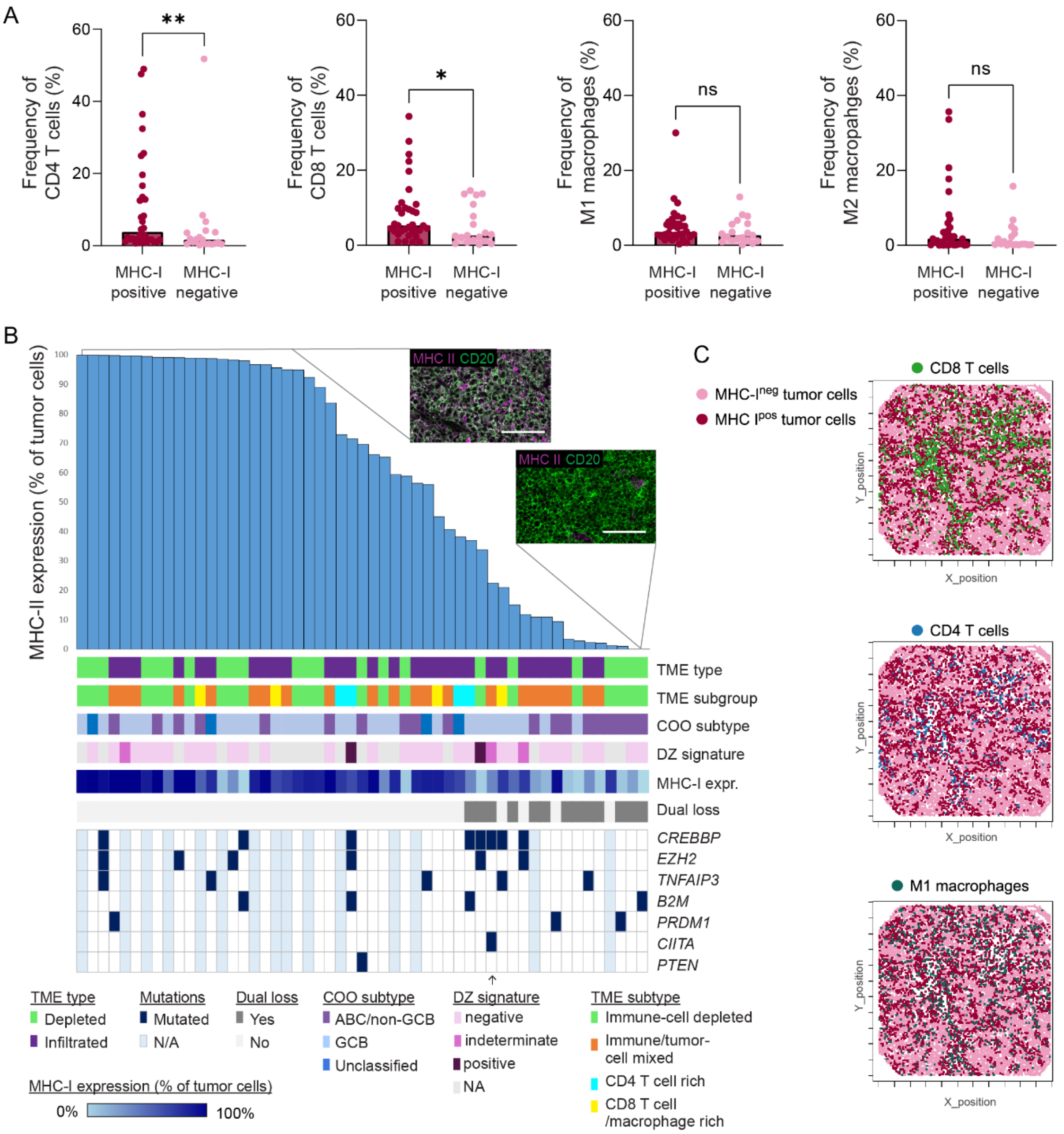


**Supplementary Fig. S7: MHC class I and II expression in DLBCL**

A) Frequency of T cells and macrophages in samples classified as MHC-I positive or negative. Bars indicate median. Mann Whitney test: * $p < 0.05$, ** $p < 0.01$. B) Samples were ordered by frequency of tumor cells expressing MHC class II. Using Youden index to identify the optimal cut-point for loss of MHC class II expression relative to TME type (96.7%) did not reveal a significant association between MHC class II expression and TME type. For further analysis, loss of MHC class II in a case was defined as less than 50% of tumor cells expressing MHC class II, resulting in 25% of cases having lost both MHC class I and II. Bars below the graph indicate TME type and subtype, COO subtype, MHC class I expression and mutational status of genes previously shown to be associated with loss of MHC class II. CD) Spatial organization of MHC class I$^{+}$ and MHC class I$^{-}$ tumor cells relative to CD8 T cells, CD4 T cells and M1 macrophages in an infiltrated case. Bars indicate median. Mann Whitney test; * $p < 0.05$, ** $p < 0.0001$.

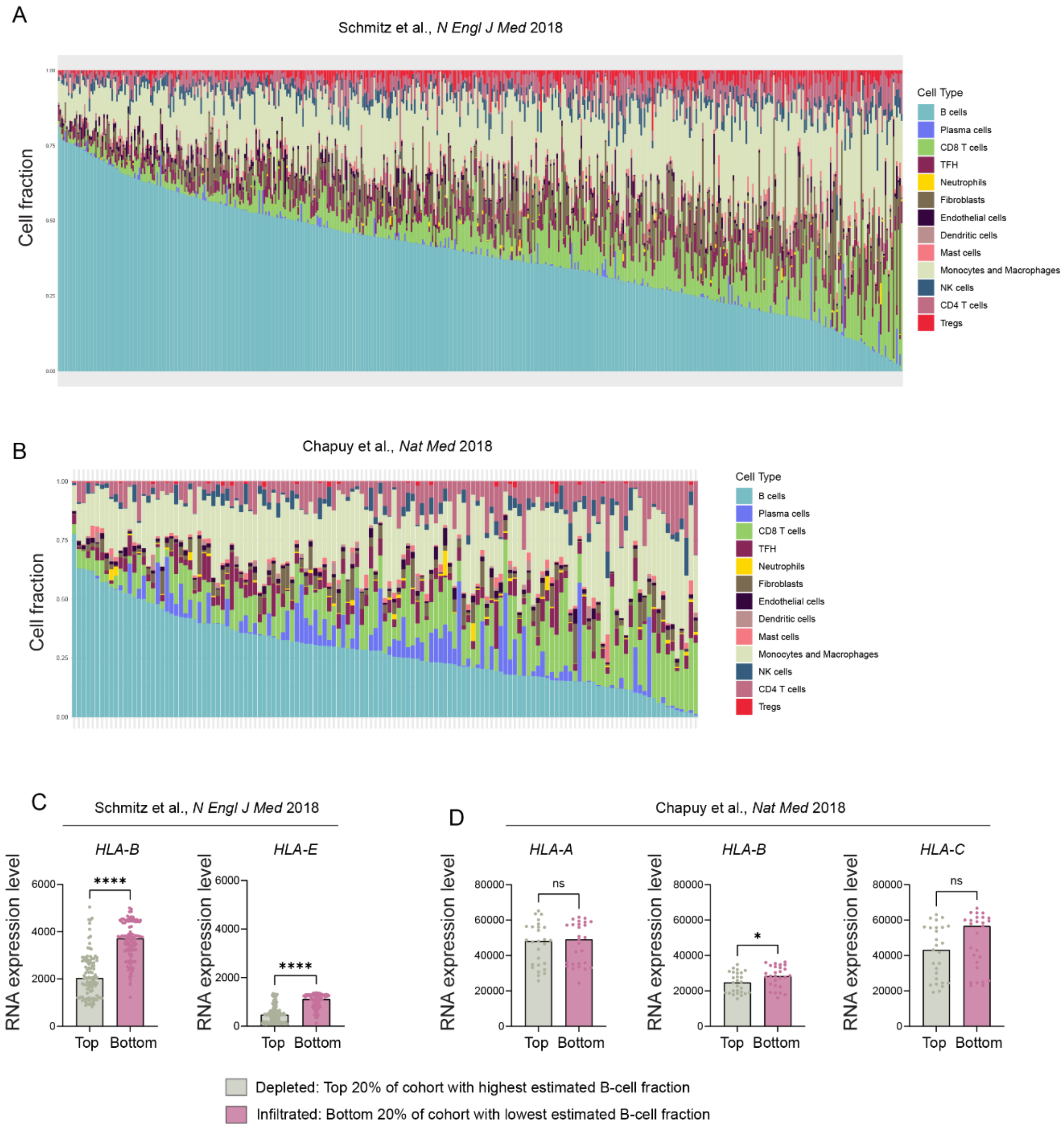


**Supplementary Fig. S8: Loss of MHC class I in B-cell rich cases validated in larger cohorts of patients**

A) CIBERSORTx were applied to the publicly available data set from Schmitz *et al.*[15] and Chapuy *et al.*[14] to deconvolute cell type frequencies from gene expression data. B) Imputed B-cell specific MHC class I gene expression (log2 transformed) was compared in cases with the 20% highest and 20% lowest frequency of B cells. Mann Whitney test; * $p < 0.05$, **** $p < 0.0001$.

### Supplementary Tables

#### Supplementary Table S1: Summary of clinical characteristics

| **Characteristics** | **All, *n* = 53** | **Depleted, *n* = 23** | **Infiltrated, *n* = 30** | ***p*-value** |
|---|---|---|---|---|
| | ***n* (%)** | ***n* (%)** | ***n* (%)** | |
| Sex | | | | |
| Male | 38 (72) | 14 (61) | 24 (80) | NS |
| Female | 15 (28) | 9 (39) | 6 (20) | |
| Age at FL diagnosis | | | | |
| Median | 54 | 54 | 55 | NS |
| Range | 36-65 | 38-65 | 36-65 | |
| Disease stage | | | | |
| II | 3 (6) | 2 (9) | 1 (3) | 0.041 |
| III | 15 (28) | 10 (43) | 5 (17) | |
| IV | 35 (66) | 11 (48) | 25 (83) | |
| Age-adjusted IPI | | | | |
| 0-1 | 2 (4) | 1 (4) | 1 (3) | NS |
| 1 | 36 (68) | 15 (65) | 21 (70) | |
| 2 | 15 (28) | 7 (30) | 8 (27) | |
| Cell of origin | | | | |
| GCB | 31 (58) | 16 (70) | 15 (50) | NS |
| Non-GCB | 22 (42) | 7 (30) | 15 (50) | |
| LDH-elevation | | | | |
| Yes | 51 (96) | 22 (96) | 29 (97) | NS |
| No | 2 (4) | 1 (4) | 1 (3) | |
| B-symptoms | | | | |
| Yes | 32 (60) | 12 (52) | 20 (67) | NS |
| No | 21 (40) | 11 (48) | 10 (33) | |
| Performance score | | | | |
| < 2 | 37 (70) | 16 (70) | 21 (70) | NS |
| $^{3}$ 2 | 16 (30) | 7 (30) | 9 (30) | |
| Bulky disease | | | | |
| Yes | 9 (17) | 5 (22) | 4 (13) | NS |
| No | 44 (83) | 18 (78) | 26 (87) | |

#### Supplementary Table S2: Antibody panel for Mass cytometry imaging

| Mass | Target | Dilution | Clone | |
|---|---|---|---|---|
| 141Pr | PAX5 | 50 | Puls Medica Device, Clone 24 | |
| 142Nd | CD34 | 50 | Leica, Clone QBEnd/10 | |
| 143Nd | HLA-II | 1600 | CST, clone LGII-612.14 | |
| 145Nd | CD56 | 100 | AH Diagnostics, Clone MRQ-42 | Excluded |
| 146Nd | CD16 | 50 | Fluidigm, EPR16784 | |
| 148Nd | ICOS | 50 | Fluidigm D1K2T | Excluded |
| 149Sm | HLA-A | 400 | Abcam, EP1395Y | |
| 150Nd | PD-L1 | 50 | Fluidigm E1L3N | |
| 151Eu | OX-40 | 50 | Fluidigm, polyclonal | Excluded |
| 152Sm | CD45 | 200 | Fluidigm CD45-2B11 | |
| 153Eu | LAG-3 | 50 | Fluidigm D2G40 | Excluded |
| 154Sm | CD366 | 400 | Fluidigm, clone D5D5R | |
| 155Gd | FOXP3 | 100 | Fluidigm 236A/E7 | Excluded |
| 156Gd | CD4 | 100 | Fluidigm EPR6855 | |
| 158Gd | FOXP1 | 50 | CST, clone D35D10 | |
| 159Tb | CD68 | 600 | Fluidigm KP1 | |
| 160Gd | TIGIT | 50 | In-house, AM15.1.2 and AM6.3.2 | Excluded |
| 161Dy | CD20 | 400 | Fludigm H1 | |
| 162Dy | CD8a | 400 | Fluidigm C8/144B | |
| 163Dy | CD163 | 50 | Leica, clone 10D6 | |
| 164Dy | CD58 | 50 | Fisher Scientific, clone 126 | |
| 165Ho | PD-1 | 100 | Fluidigm EPR4877(2) | Excluded |
| 166Er | CD45RA | 10000 | Fluidigm HI100 | |
| 167Er | GrzB | 200 | Fluidigm EPR20129-217 | |
| 168Er | Ki-67 | 100 | Fluidigm B56 | |
| 169Tm | Collagen | 400 | Fluidigm Polyclonal | |
| 170Er | CD3 | 100 | Fluidigm Polyclonal | |
| 171Yb | Hist H3 | 2000 | Fluidigm D1H2 | |
| 173Yb | CD45RO | 800 | Fluidigm UCHL1 | |
| 174Yb | HLA-B | 200 | Nordic Biosite, polyclonal | |
| 175Lu | CD54 | 50 | CST, clone E3Q9N | |
